# MutateX: an automated pipeline for *in-silico* saturation mutagenesis of protein structures and structural ensembles

**DOI:** 10.1101/824938

**Authors:** Matteo Tiberti, Thilde Terkelsen, Tycho Canter Cremers, Miriam Di Marco, Isabelle da Piedade, Emiliano Maiani, Elena Papaleo

## Abstract

Mutations resulting in amino acid substitution influence the stability of proteins along with their binding to other biomolecules. A molecular understanding of the effects induced by protein mutations are both of biotechnological and medical relevance. The availability of empirical free energy functions that quickly estimate the free energy change upon mutation (ΔΔG) can be exploited for systematic screenings of proteins and protein complexes. Indeed, *in silico* saturation mutagenesis can guide the design of new experiments or rationalize the consequences of already-known mutations at the atomic level. Often software such as FoldX, while fast and reliable, lack the necessary automation features to make them useful in high-throughput scenarios. Here we introduce MutateX, a software which aims to automate the prediction of ΔΔGs associated with the systematic mutation of each available residue within a protein or protein complex to all other possible residue types, by employing the FoldX energy function. MutateX also supports ΔΔG calculations over protein ensembles and the estimation of the changes in free energy upon post-translational modifications. At the heart of MutateX lies an automated pipeline engine that handles input preparation, performs parallel runs with FoldX and outputs publication-ready figures. We here illustrate the MutateX protocol applied to the study of the mutational landscape of cancer-related proteins, industrial enzymes and protein-protein interfaces. The results of the high-throughput scan provided by our tools could help in different applications, such as the analysis of disease-associated mutations, or in the design of protein variants for experimental studies or industrial applications. MutateX is a collection of Python tools that relies on Open Source libraries and requires the FoldX software to be installed beforehand. It is available free of charge and under the GNU General Public License from https://github.com/ELELAB/mutatex.

## Introduction

Advances in the field of proteomics are now providing a huge amount of data on protein-protein interaction or post-translationally modified proteins ^1–5^ that benefit of structural studies to make a rational out of them ^6^. On other hand, genomic initiatives allow to identify missense mutations^7,8^ in the coding region of genes that need to be understood at the structural and functional level, along with regarding the impact of mutations on protein stability ^9,10^.

Indeed, single amino acid substitutions (i.e. mutations) or post-translational modifications (PTMs) in proteins may result in significant changes in protein structural stability and intermolecular interactions, impacting on protein activity, function and cellular signaling. An accurate and systematic prediction of changes in stability and binding upon mutations or PTMs is thus crucial to understand protein variants at the molecular level ^11–13^. Moreover, the possibility to predict mutational effects also has a strong impact in protein engineering for biotechnological and industrial applications ^14,15^. It has been also shown that disease-causing mutations are characterized by changes in structural stability or binding affinity ^16–18^. Most prediction tools are designed to work on a single conformation at a time, however proteins are not static entities and they can attain multiple states in solutions that are tightly related to function ^19,20^. Some proteins are conformationally heterogeneous and undergo a myriad of structural changes, including sparsely populated states that are stabilized upon interaction with a biological partner ^21,22^. It is thus important, for a proper understanding of the impact of mutations and PTMs, to account for the conformational ensembles of proteins. Protein ensembles can be obtained by experimental techniques such as NMR ^23,24^, simulations ^25–27^ or an integration of the two ^28–30^.

Many diverse methods have been developed to estimate stability and binding free energy changes upon amino acidic substitutions from a structure, many of which extensively summarized in a recent review article. ^31^ Given their widespread use and importance, synthetic benchmarks of them have been performed over time, along with comparisons with experimental data. These structure-based methods were in general considered to be moderately accurate in terms of classification between stabilizing, neutral or destabilizing mutations ^32,33^ and in terms of correlation between experimental and computed ΔΔG values ^34,35^. Among these methods, FoldX ^36^, has been ranked among the best-performing methods ^37^ and has been applied with success to many cases of study and biological questions. Experimental validation of FoldX predictions, along with comparison to existing experimental data, also support the notion that this tool is valuable for mutation classification, design and prediction ^31,37–41^.

The FoldX method predicts changes in free energy of folding upon mutation as the difference between the estimated free energy of folding of the mutant and the reference wild type variants. This is done by using an empirical free energy function, which includes terms for Van der Waals interactions, solvation free energies, water bridges, hydrogen bonding, electrostatics and entropy changes upon folding for main-chain and side-chains. Some of these terms are weighted by coefficient, which was determined using a fitting procedure over a database of free energy differences from single-point mutants, meaning that FoldX free energies are not necessarily physical – however, the calculated differences are comparable to those identified experimentally. The energy function is implemented in a closed-source commercial software suite, FoldX, which is available for free for academic use and is able to calculate the free energies estimate and to energy minimize and mutate input protein structures. The suggested FoldX mutation protocol includes a highly recommended Repair step, in which an energy optimization is performed that removes interatomic clashes or otherwise changes the conformation of residues with particular bad energy in the input structure. Next a mutation step is performed (*BuildModel*) in which the model of the mutant variant and corresponding wild-type models are generated, and finally the respective energies are calculated and subtracted to obtain the final difference in free energy upon mutation. FoldX is also able to estimate the free energy of interaction between pairs of molecules in the structure through the *AnalyseComplex* command, in which the energy of interaction is simply calculated as the difference between the energy of the isolated molecules with that of the complex. FoldX supports the twenty canonical amino-acids, few post-translationally modified ones and other chemical moieties such as nucleic acids and metal ions.

The accuracy of FoldX has been independently investigated in literature, with a standard deviation reported in the range of 1.0-1.78 kcal/mol between the FoldX ΔΔGs and experimental data, and an average accuracy of 0.69 ^37^. Among the methods tested for systematic bias, it was found to be among the top five least biased methods ^42^, with tendency towards destabilizing mutations. The bias itself can be alleviated through a backbone relaxation step ^43^.

Interestingly, FoldX has been found to be quite sensitive to the input structure and using high-quality crystal structure for ΔΔG determination has been suggested. In this regard, a modified protocol in which ensemble averages of free energies predictions calculated on conformers from molecular dynamics simulations has been proposed ^41^, with the observation that, while predictions performed on the single MD conformation had quite poor correlation with experimental ΔΔG, using ensemble average predictions restored the correlation and reduced the spread of the ΔΔG values. This indicates that ensemble averaging in FoldX constitutes a viable line of research that could improve the results obtainable by the software.

The aforementioned energy functions, and FoldX in the first place, are, in general, promising approaches since they, in principle, allow for a fast but yet quantitative estimate of the changes in stability and at the interaction interface for other biological partners of all possible mutations in a protein structure. Nevertheless, elegant computational solutions to design flexible and comprehensive pipelines to carry out *in*-*silico* deep mutational scans in a high-throughput manner are still missing, along with the extension of these functions to a systematic study of protein ensembles taking advantage of parallelization on modern multi-core hardware.

## Results

### Design and Implementation

#### Overview on MutateX

MutateX includes a wrapper in Python for the FoldX suite accompanied by plotting and post-processing Python scripts. The main program implements an *in*-*silico* saturation mutagenesis protocol for proteins and protein complexes that uses the popular FoldX program to predict the effect of mutations. It allows to calculate the free energy change associated with the systematic substitution of each protein residue to any of the natural amino acids or post-translational modified residues, such as phospho-residues. MutateX automates and streamlines the mutation process, making it straightforward to run a full mutational scan even for several conformers of the same protein. From a single three-dimensional (3D) protein structure, or ensemble of structures, MutateX provides the researcher with textual and graphical representations of the mutational data to explore and make sense of the huge amount of information derived from such an exhaustive scan. This overview is possible through the use of powerful visualization tools, which produce publication-ready figures.

At the heart of the MutateX software lies a pipeline engine that handles input preparation, parallel runs of available programs able to estimate the free-energy, coupled with data gathering and visualization tools. Currently, the supported software for the calculation of free energies is the FoldX Suite, which employs the empirical FoldX energy function ^36,44^ to determine the energetic effect of point mutations, along with the binding energies associated with the formation of multimeric protein complexes. MutateX is compatible with FoldX Suite version 4 and the recently relased version 5 ^45^.

#### The MutateX protocol

In its simplest application, the MutateX main wrapper requires as an input a single file with the coordinates of a protein 3D structure in the PDB format. Moreover, the user should provide the FoldX running input files, for which templated are included by default in the MutateX distribution. The templates for the different calculations are used by FoldX for the *Repair, BuildModel* and (optionally) *AnalyseComplex* FoldX commands, as detailed in the User Guide. Advanced users are invited to create their own templates or modify the existing ones as they see fit.

The MutateX software implements the protocol summarized in **Fig. 1.** First, the main MutateX program parses the input PDB files and saves one file per model in the PDB format. As a second step MutateX prepares and performs a *Repair* FoldX run on each model. For each repaired model, Mutatex generates all the directories and files that are required to perform the mutational scan, including structure files, run files for FoldX, and lists of mutations. All substitutions on a given site are treated as a single independent FoldX run, making it possible to run several instances of the FoldX software at the same time, which is necessary to take advantage of modern multi-core processors and effectively reduce the execution time of the whole pipeline. MutateX takes care of scheduling the necessary FoldX runs, keeping track of their status and checking the output files. During the preparatory step, the software automatically recognizes if two or more identical protein chains are present in the input structure files and prepare the mutation runs taking this into consideration, by mutating all the chains having the same sequence at the same time. It also supports setting the number of runs per mutation, so that cases in which more local structural variability is expected can be taken into account. Once all the mutation runs have been performed, MutateX reads the output of the FoldX runs and summarizes the estimated average free energy differences, the associated standard deviations and minimum and maximum values in separate text files for each residue. The MutateX processing tools can then be used on these files to obtain representations of the effects of the mutations, such as probability density plots of ΔΔG values calculated with Kernel density estimation (*ddg2density*), heatmaps (*ddg2heatmap*), histograms relative to single mutation sites (*ddg2histo*), logo plots (*ddg2logo*), per-site distributions of ΔΔGs in different representations (*ddg2distribution*). When plotting, the user can specify custom labels by filling in a template generated using the *pdb2labels* tool. The *ddg2dg* tool can be used to apply systematic linear corrections to the calculated ΔΔG values. The dataset can be converted to Microsoft Excel-compatible files (*ddg2excel*) or PDB files with relevant information about the mutagenesis results stored in the B-factor field (*ddg2pdb*). These files can be further visualized and processed using protein structure viewers, such as PyMOL (https://pymol.org). The same data can also be written as the diagonal of a residue matrix file, compatible with the xPyder plugin ^46^. The user can also generate ad-hoc reports on specific residues (ddg2summary). Most of these tools support extensive plotting options, the most important of which is the range of ΔΔG values that are to be considered. In fact, according to our experience, the ΔΔGs predicted by FoldX can go up few tens kcal/mol. Therefore, if the entire range of predicted ΔΔGs is considered, identifying smaller but still meaningful differences becomes difficult. We suggest to limit the range of ΔΔG values between −3.0 and 5.0 kcal/mol, as this is the range in which most mutations were experimentally found (see Results for details). ΔΔG values below or above this are likely to be overestimated by the method.

**Figure 1:**
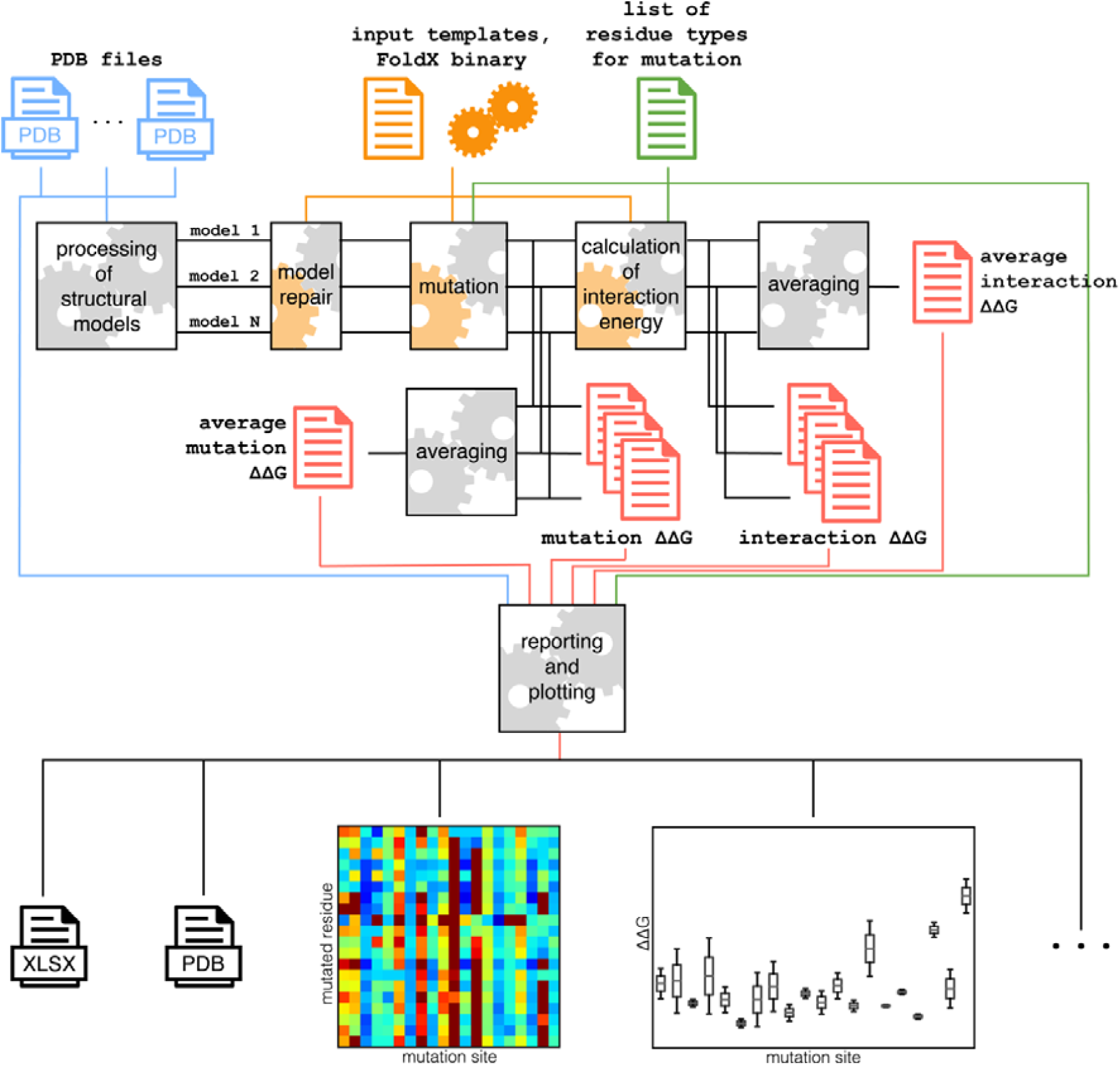
Scheme of the workflow implemented in MutateX, as described in the Methods section.

Self-mutations, i.e. those mutations that replace a residue with one of the same type, can also be performed independently respect to the rest of the protocol and are especially useful as a sanity check of the method for every single mutation site, in a given structure. In fact, the free energy difference calculated when mutating a residue to itself acts as a lower bound of the prediction error, as we would normally expect ΔΔG close to 0 for such a mutation.

MutateX allows to customize some aspects of the calculations. For instance, it is possible to provide the list of target residue types that should be considered in the scan, as well as to include multiple models from one or more PDB files. Indeed, MutateX supports the saturation scan and the free energy calculations over structural ensembles, an approach that in principle might be able to mitigate the inaccuracies associated with the limited conformational variability that FoldX allows for during modelling of the mutant variants. The software is also able to consider the simultaneous introduction of each mutation on more than one chain at the same time in the case homo-dimers or homo-multimers are present in the PDB file. Finally, MutateX is designed to handle, calculate and plot the changes of free energies of binding between complexes upon mutation, both considering homo-multimers and hetero-multimers.

#### Performance

The vast majority of run time for MutateX scan is taken by the runs of the ΔΔG estimation engine, as the operations of preparation and post-processing performed by MutateX are computationally inexpensive. Therefore, the performance of MutateX is strictly tied to that of the underlying mutation engine – FoldX in this case. MutateX is able to run a complete mutational scan of a 300-residues protein to all the 20 natural variants in less than a day using 4 cores for parallel execution, which can routinely be done on a desktop-class machine (**Fig. 2**).

**Figure 2.**
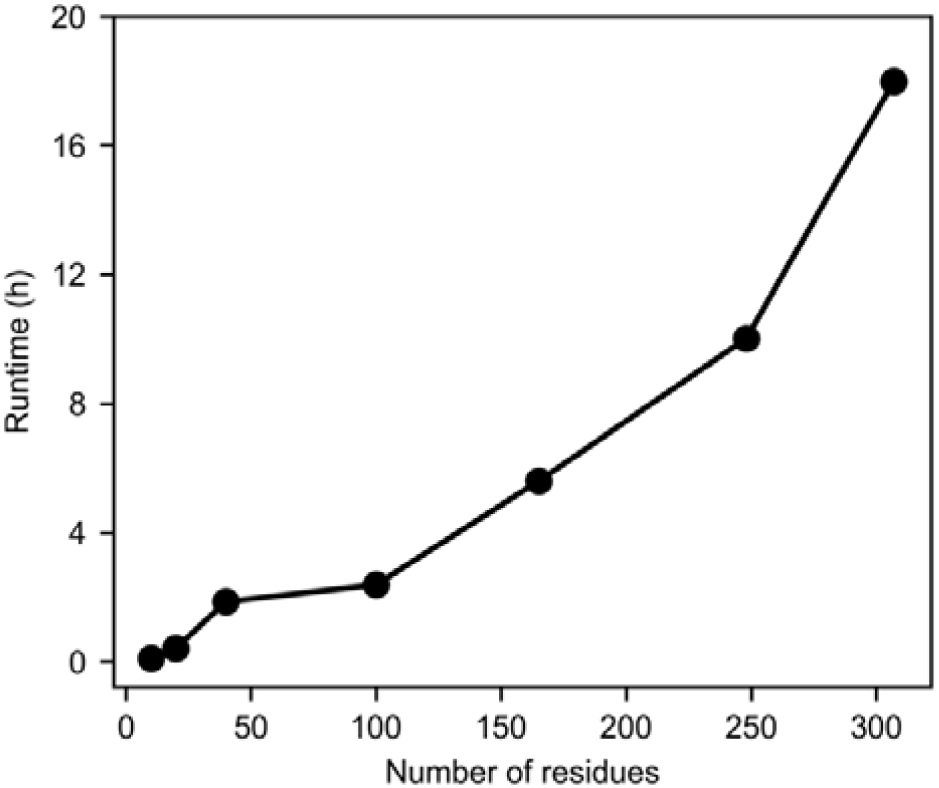
Scaling behavior of MutateX with increasing number of residues. These run times have been obtained using MutateX in combination with FoldX Suite version 4.

It should be noted that, as the FoldX runs performed by MutateX are completely independent on one another, the scaling capacity of MutateX itself is linear with the number of cores, peaking when the number of cores is equal to the number of residues to be mutated.

## Results

### Selection of a cutoff for truncation of ΔΔGs values for plotting purposes

The predicted FoldX ΔΔGs can reach up few tens of kcal/mol in some cases, making visualizing a full mutational scan challenging due to scale effects, for instance when using heatmaps. In fact, using large scales would mask smaller but meaningful differences difficult to read, giving more importance to very large energy differences that are more likely to be over-estimations. We therefore sought to identify a range of free energy differences in which small differences are to be considered meaningful, while smaller or larger values than that can be flattened out as largely destabilizing or stabilizing. To this extent, we have used the ProTherm database to check how experimentally-derived ΔΔG values are distributed, after filtering out those values for which an energy unit was not specified in the database. **Fig. 3** shows that most mutations fall within the −3.0 to 5.0 kcal/mol range, which can be used as a cutoff for truncation of DDG values when visualizing MutateX mutation results, which correspond roughly to the second and 98th percentile respectively. We therefore recommend using these or more extreme values as range limits for visualization and plotting purposes.

**Figure 3.**
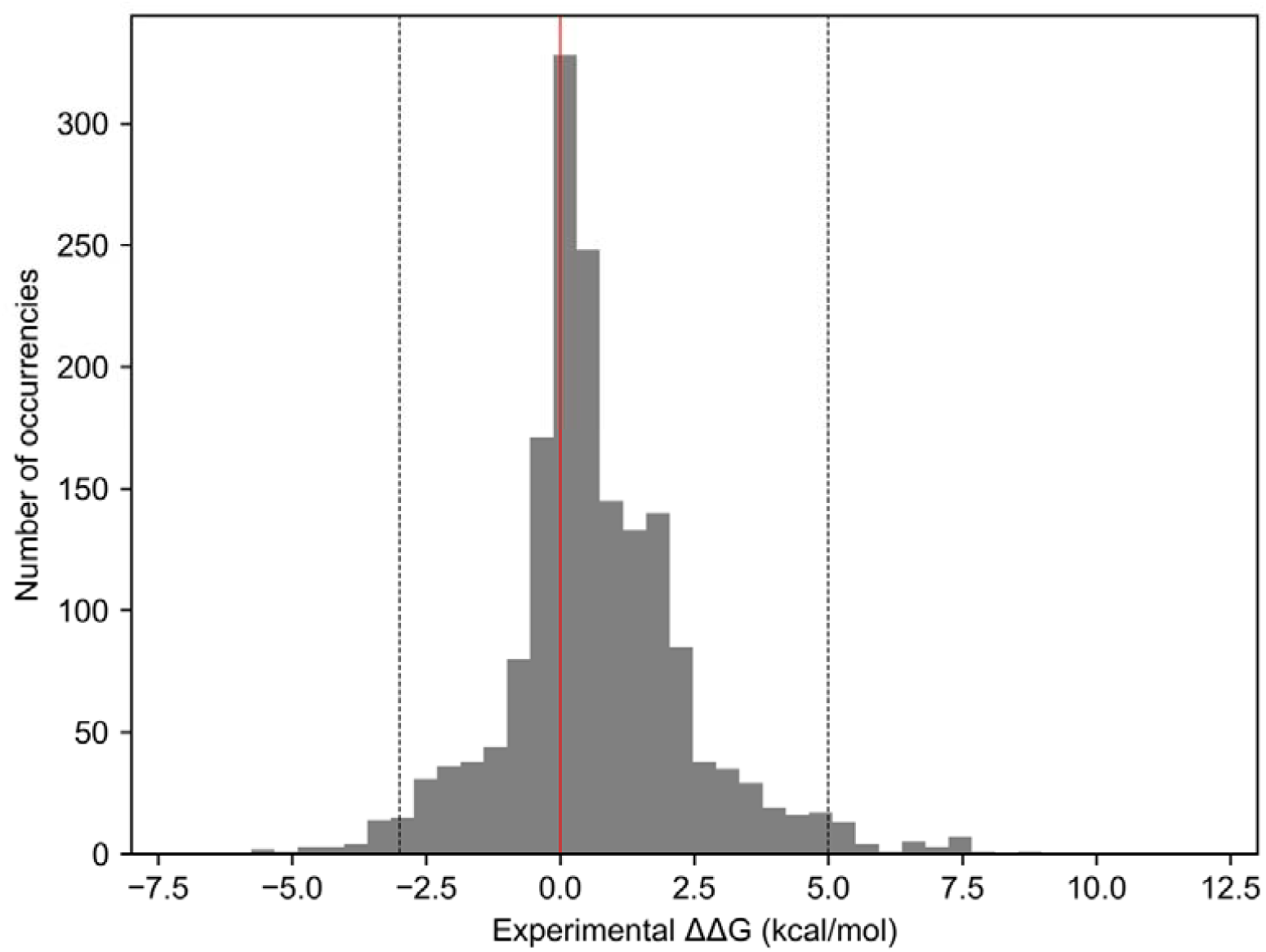
Distribution of experimental ΔΔG values, upon sign change, as found in the ProTherm database ^47^for which a unit was specified.

### Examples of use

We show as examples of usage for MutateX the prediction of: i) protein-destabilizing or stabilizing mutations through a full mutational scan of the cancer-related protein, Phosphatase and tensin homolog (PTEN), ii) an example of protein-protein interaction with the tetrameric structure of the p53 DNA binding domain, iii) an example of protein-peptide interaction, including the estimate of effects of phosphorylation, with the mutational scan of complexes between SH2 and phospho-peptides.

### Saturation mutation scan of the cancer-related protein PTEN: relationship between protein stability and disease

PTEN is a tumor suppressor with homology to tyrosine phosphatases and the cytoskeletal protein tensin ^48^. The PTEN monomer consists of two major functional domains, i.e. the phosphatase domain (PTP, residues 14-185) and the C2 domain (residues 190-285). PTEN is known to be involved in regulation of cellular growth processes and suppresses cell migration and invasion ^49^. Mutations in PTEN have been identified in a variety of human cancers ^50^. Most reported missense mutations of PTEN are located within the exon encoding the PTP domain.

PTEN has been recently used as a case study, in which other methods have been used to predict changes in free energy upon mutations and then compared to multiplexed experiments ^51^. The authors found that the loss of PTEN stability was a driver factor for disease-causing mutations with 60% of the pathogenic variants causing the loss of function because the protein is destabilized and degraded. PTEN is thus a useful case study to illustrate the possible application of the scan for changes in stability with the MutateX protocol. We performed a saturation mutagenesis scan of PTEN, aiming at identifying the most damaging mutations and relate them with available mutational data from publicly available cancer samples.

We employed the most complete structure of PTEN available at the best of our knowledge (PDB ID 1D5R ^52^). Notably, the PDB entry lacks atomic coordinates for residues 286-309 in a disordered region – however, given the local sampling carried out by FoldX, the missing residues are likely to affect only residues in the proximity of the loop. We thus did not model the missing region since this calculation serves only for illustrative purposes.

The results from the scan were visualized exploiting different post-processing tools available within MutateX. We first analyzed the outcome of the scan using heatmaps (produced by the *ddg2heatmap* tool) and boxplots (*ddg2distribution*), which provide a comprehensive overview of the free energy changes upon mutations along the protein sequence. In particular, the *ddg2distribution* tool has been designed to produce different representations of the obtained ΔΔGs on a per-residue distribution basis. The heatmap and box plots shown in **Fig. 4A and 4B** depict the change of ΔΔGs in the region 64-113 of PTEN, corresponding to the middle part of the phosphatase domain. The results may be classified into the following scenarios:

i. the ΔΔG is not influenced by substitution of the wild-type amino acid. An overall lack of change in ΔΔGs at a certain site indicates that the wild-type residue is not essential for protein stability. This is often the case when residues that are located at the protein surface and lack interactions with the neighboring residues, or for those residues located in flexible regions (such as the residue D77).
ii. The ΔΔG increases upon mutations of the wild-type residue for a certain subset of other residues. This subset is likely to have something in common, e.g. residues are hydrophobic, hydrophilic, charged, small/large or have an aromatic side chain. Substitutions of a small uncharged residue for another is unlikely to alter protein stability, whereas substitution to a large aromatic side chain could have a marked effect (such as in the case of N69).
iii. The ΔΔG increases regardless of what the wild-type amino acid is mutated to, indicating that the native residue may play an essential/specific role in the preservation of the protein structure (such as the example of P95 and P96).
iv. The ΔΔG decreases when the wild-type residue is mutated to one or a subset of other residues – e.g. the substitutions result in a more stable protein variant. An observed decrease in ΔΔG is expected to be significantly more modest is size and observed less often as protein structure/stability is evolutionary optimized (on example is the substitution of residue Q with Y or F at position 87).

**Figure 4.**
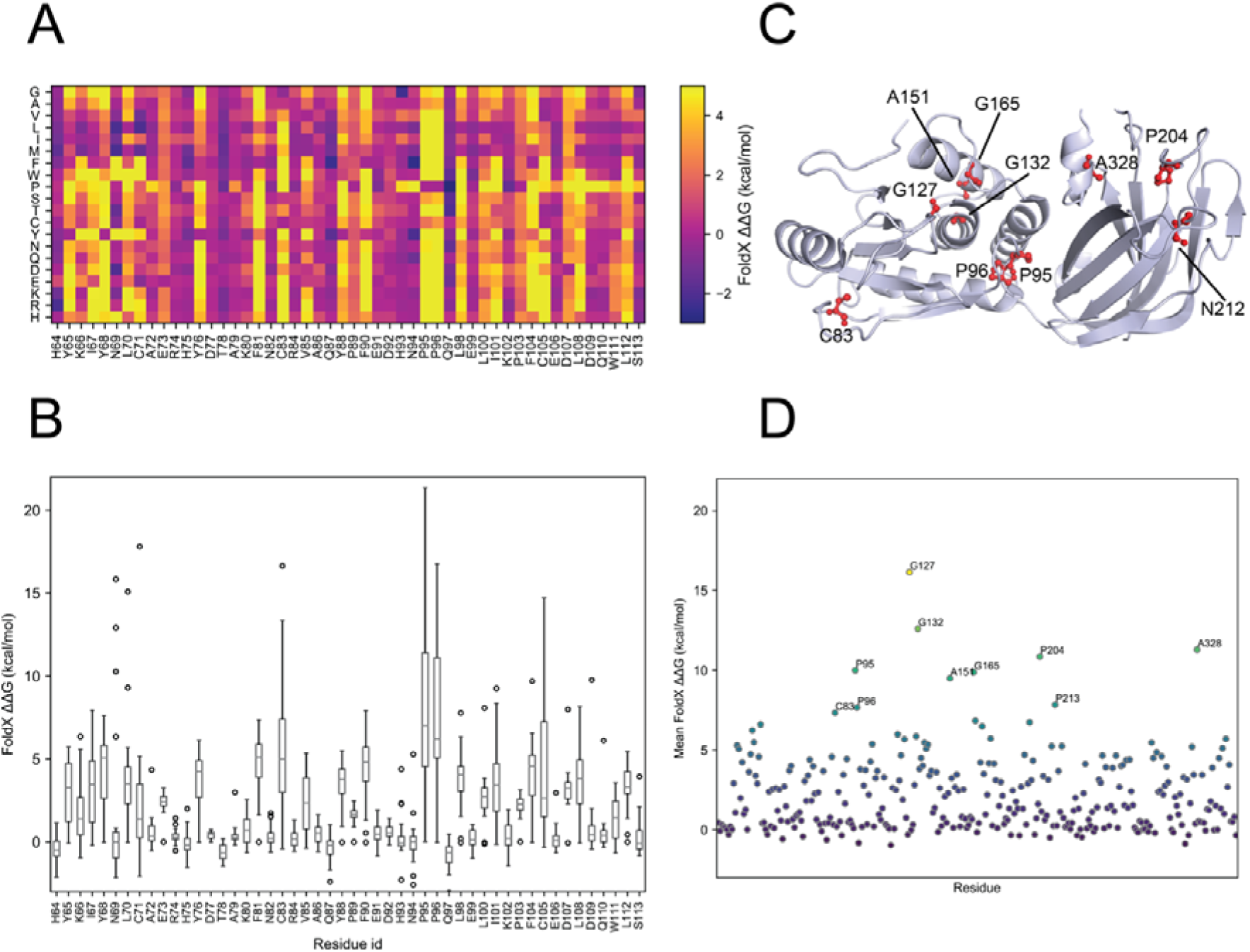
Saturation mutagenesis scan of the tumor suppressor PTEN by MutateX. The heatmap (A) and box plot (B) of the mutational scan corresponding to the residues 64-113 is shown as an example. It includes the folding ΔΔGs estimated for the mutation of every mutation site (x axis) to every natural amino-acid (y axis) using FoldX. The values of free energy difference have been limited to the range -3-5 kcal/mol, meaning that values above 5 or below −3 kcal/mol are shown as the extremes of the range. In panel D, the average ΔΔG for the mutation of each residue to all the 20 natural residue types is shown; then the ten sites with largest average ΔΔG are labeled. The same mutations are shown on the structure of PTEN in panel D, produced with PyMOL.

To determine which of the 351 PTEN amino acids were most important for protein stability, we have used ddg2distribution to plot the per-site average ΔΔG over all of the residues of PTEN. We have further annotated with residue names the ten positions with the largest average ΔΔG (**Fig. 4C and 4D**).

The residues that on average have a larger effect on stability upon mutation for PTEN stability were found to be position 83, 95, 96, 127, 132, 151, 165, 204, 213 and 328. The top 10 residue are highlighted with dots/sticks on the 3D structure of the protein reported in **Fig. 4C.** The mutation predicted to be most sensitive to substitution is the glycine residue at position 127, which is located in a loop right before the third α-helix in PTEN. Glycine residues located in disordered regions such as loops provide a certain degree of flexibility, which is important for the proper functional breathing motions of the protein ^26^. It should also be noted that, since FoldX does not relax the protein backbone upon mutation, replacing a residue without side-chain with larger ones can result in steric clashes that artificially increase the obtained ΔΔG values.

The MutateX full mutational scan of PTEN also allowed us a better understanding of the effects induced by disease-related mutations. As PTEN functions as a tumor suppressor in many cancer types we extracted 1116 missense cancer-associated mutations of PTEN using the cBioPortal database ^8^ (August 2019) -a comprehensive collection of cancer mutations identified by genomic initiative of high-throughput profiling of samples from cancer patients. Intriguingly, eight out of ten of the top ten candidates identified in the MutateX scan were annotated in Cbioportal and are reported in **Table 1.** In particular, mutations of G127, G132 and G165 were among the most frequent PTEN mutations in cancer patients.

**Table 1.** Residues with largest average ΔΔG annotated with information from cBoPortal.

| residue position | WT amino acid | Annotated mutant amino acid | Number of missense annotations | Cancer type | Location within PTEN structure |
| --- | --- | --- | --- | --- | --- |
| 83 | C | NA | NA | NA | Phosphatase domain |
| 95 | P | S, L | 5 | Glioblastoma Multiforme, Uterine Endometrioid Carcinoma, Colorectal Adenocarcinoma, Mixed Germ Cell Tumor | Phosphatase domain |
| 96 | P | A, L, S, T | 4 | Uterine Endometrioid Carcinoma, Chromophobe Renal Cell Carcinoma, Glioblastoma Multiforme, Uterine Endometrioid Carcinoma | Phosphatase domain |
| 127 | G | E, R, V | 15 | Lung Adenocarcinoma, Glioblastoma Multiforme, Hepatocellular Carcinoma, Cutaneous Melanoma, Uterine Endometrioid Carcinoma, Colorectal Adenocarcinoma, Ampullary Carcinoma, Teratoma, Uterine Clear Cell Carcinoma, Uterine Serous Carcinom | Phosphatase domain |
| 132 | G | A, C, D, S, V | 23 | Solitary Fibrous Tumor, Uterine Carcinosarcoma, Glioblastoma Multiforme, Cutaneous Melanoma, Colorectal Adenocarcinoma, Lung Adenocarcinoma, Astrocytoma, Uterine Endometrioid Carcinoma, Esophageal Squamous Cell Carcinoma, Medulloblastoma, Prostate Adenocarcinoma, Rectal Adenocarcino | Phosphatase domain |
| 151 | A | G, T, D, V | 4 | Breast Invasive Lobular Carcinoma, Colorectal Adenocarcinoma, Uterine Carcinosarcoma, Glioblastoma Multiforme | Phosphatase domain |
| 165 | G | E, R | 15 | Small Cell Lung Cancer, Renal Non-Clear Cell Carcinoma, Medulloblastoma, Cutaneous Melanoma, Glioblastoma Multiforme, Rectal Adenocarcinoma, Uterine Endometrioid Carcinoma | Phosphatase domain |
| 204 | P | L, R, S | 7 | Melanoma, Glioblastoma Multiforme, Uterine Endometrioid Carcinoma, Uterine Carcinosarcoma, Colorectal Adenocarcinoma, Cutaneous Melanoma | C2 domain |
| 213 | P | R | 1 | Prostate Adenocarcinoma | C2 domain |
| 328 | A | NA | NA | NA | C2 domain |

### Saturation mutagenesis to identify hotspot residues at the binding interface for tetramerization of p53

The tumor suppressor p53 is known as the guardian of the human genome. It is a stress-response transcription factor that is involved in different cellular functions such as cell cycle arrest, senescence and apoptosis ^53^. Mutations in the p53 gene occur at high frequency in a large variety of cancers ^54–56^. p53 exerts its function as a transcription factor by binding to DNA through its central folded DNA-binding domain (DBD, residues 94-292), which is also able to bind other proteins and acts as a signal integration hub ^53,57,58^. The activity of p53 depends on its conformation: p53 is active in a tetrameric (dimer of dimers) conformation which is required for its ability to bind with high affinity to DNA and interact with various proteins ^53^. Formation of the tetramer happens through a specialized tetramerization (TET) domain; nonetheless, the DBD of the four p53 units in the tetramer still interact with one another and the binding is stabilized by protein-protein interactions ^59^. Many of the cancer-related p53 mutations are located in the DBD and have destabilizing effects, but there are also mutations which do not fall within this class. To fully understand the effects of mutations on p53 and their roles in cancer, it is thus essential to identify hotspot mutations that could also induce structural changes at the binding interface for tetramerization of p53. As an example, we used MutateX to study the effects of mutations on the binding interfaces between DBDs in the p53 tetramer. We performed a saturation mutagenesis scan of p53 tetramer, mutating each residue to all the 20 natural amino acids, using the crystal structure of the p53 DBD as a self-assembled tetramer (PDB ID 3KMD) ^60^ and calculated the differences in binding energies upon mutation. As the structure contains four protein chains with identical sequence, MutateX automatically recognized it as a tetramer and therefore performed the same mutation of each chain for every run. The software then calculated differences in binding energy considering every possible combination of protein chains (A-B, B-C, C-D, A-D, A-C, B-D). The p53 dimer of dimer has a planar structure shaped as a parallelogram, with no contacts between monomers A-C and B-D. Interface A-B connects two p53 monomers into a dimer (dimer interface), while interface A-D connects two dimers into the tetramer (dimer-dimer interface). We here considered only differences in free energy of binding between pairs A-B and A-D, as the other two interfaces are identical to these. We first plotted a heatmap of binding ΔΔGs using *ddg2heatmap*, which provides a comprehensive visual representation of the free energy changes, limiting the plotted values between −3.0 and 5 kcal/mol **(Fig. 5A and 5C)**. We’ve also plotted the per-site ΔΔG distributions using ddg2distribution, obtaining either a violin or a stem plot **(Fig. 5B and 5D)**. Among the tested mutations, those that significantly change the free energy of interaction between chains A and B are in the regions C176-R181 and S241-C245 as shown in **Fig. 5A.** These correspond indeed with the dimerization interface of the two p53 chains A and B, which includes part of L2, around its short alpha-helix and L3, close to where the zinc-binding site is located. Some key residues that were previously identified as important for forming the binding interface, such as P177, H178, E180, M243 and G244 and surrounding residues are those that show the highest binding ΔΔG increase when mutated. For instance, mutations of residue C176 to large aromatic residue (F, W, Y) or charged ones (K, R, H) affect binding significantly, while in position P177 S, T, G and R are the most damaging mutations. Replacing key residues on the other side of the binding interface is predicted to be damaging in general, given the high ΔΔG we MutateX computed for mutations at M243 and G244. All the substitutions for M243 and G244 are predicted to result in weakened binding, with L, I and M being the most tolerated substitutions (binding ΔΔG < 1.0 kcal/mol).

**Figure 5.**
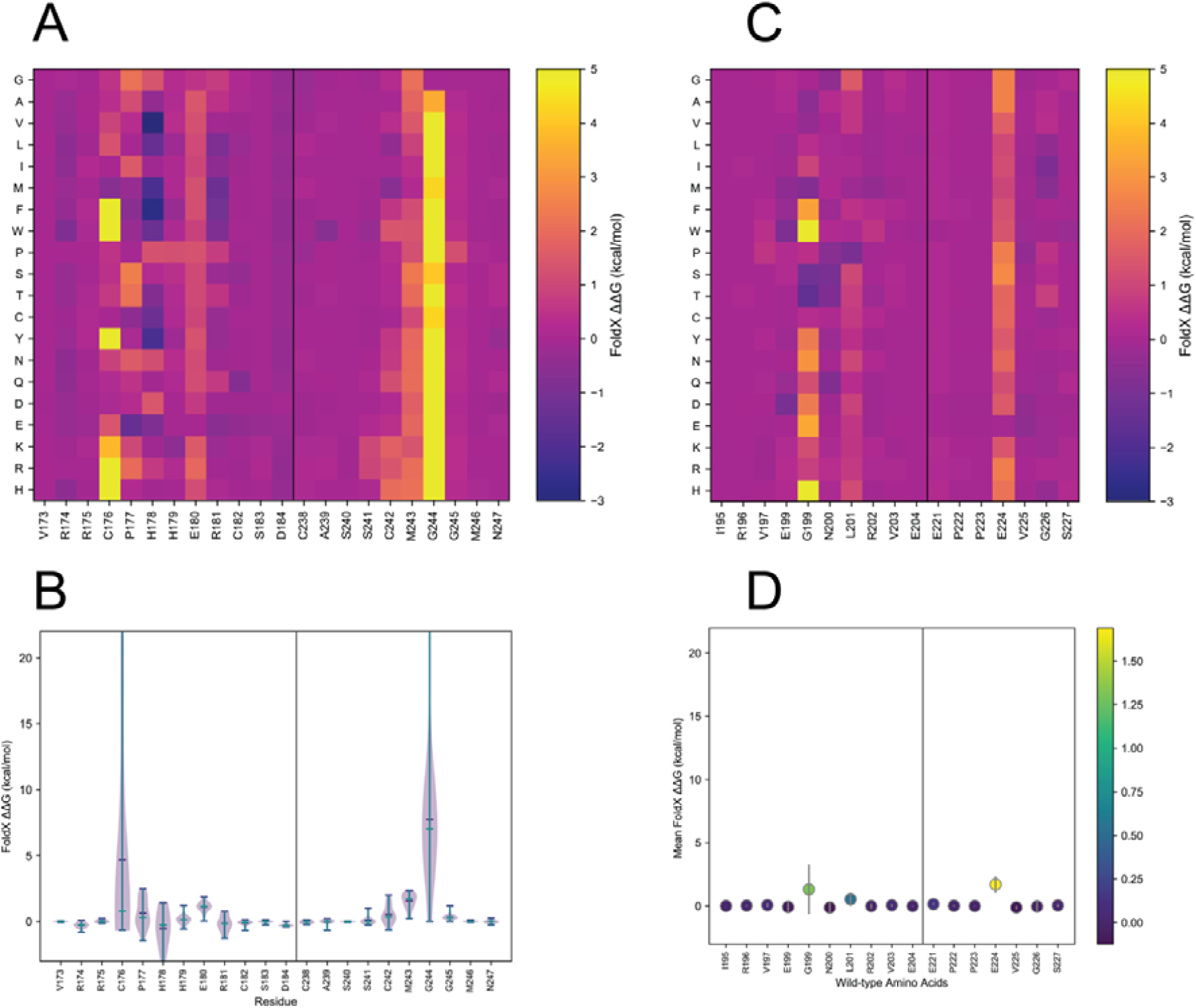
Saturation mutagenesis for tetramerization of p53. A. Heatmaps of interaction ΔΔG for p53 generated using MutateX. Only the residues that take part in the interaction between the different chains are shown. On the left column, the difference interaction energies are between chains A and B, while on the right they are for chains A and D. Heatmaps for both cases have been generated using the *ddg2heatmap* tool, white a violin plot and a stem plot have been generated using *ddg2distribution*. Each point in the stem plot shows the average difference in interaction energy of that mutation site on the 20 reisdue types its been mutated to, with the standard deviation show as a vertical bar.

Next, we analyzed the interaction surface between chains A and D, which corresponds to the dimer-dimer interface. The interface is broader and includes two patches of residues, formed by loops connecting the beta strands as well as the DBD N-terminal tail and loop 2. One patch of this interface (patch I) is described as formed by L93, S94, S95, S166, Q167, T170, and F212 of monomer A and L201, and T140, E198, G199, M200, R202, H233 of monomer D. Of these residues, our scan identifies L93, Q167, G199, L201 and H233 as those harboring the most destabilizing mutations. In particular, G199 is deeply buried within patch 1 and faces T170 directly; its mutation to bulky amino acids (such as F, W, Y, H, but also D and E) is found to be very destabilizing. It should be noted that FoldX does not rearrange the protein backbone after side-chain replacement and remodeling, meaning the mutation might be predicted as more destabilizing than it would be if the whole structure could be fully relaxed. Patch 2 is smaller and composed of residue Q100 and K101 on monomer A and E224 and V225 on chain D. Of these potential mutation sites, E224 is predicted to be the most sensitive by far, with most mutations predicted to be destabilizing.

### Saturation mutagenesis of protein-peptide complexes: identifying binding specificities of SH2 domains

Protein post-translational modifications (PTMs) play a critical role in cellular transduction. Phosphorylation is the most well-characterized PTM, and consists in the transfer of the terminal phosphate group from adenosine triphosphate (ATP) to the hydroxyl group of a serine, threonine or tyrosine residue. Protein kinases and phosphatases control the levels of protein phosphorylation, regulating several processes such as proliferation, DNA repair, programmed cell death, differentiation, and immune responses ^61^. Indeed, the physiological phosphorylation pattern is frequently altered in pathological conditions such as cancer ^62^. Phosphorylation of short linear motifs creates docking sites recognized by binding domains, such as Src homology 2 (SH2) and others ^63^; SH2 domains specifically bind pTyr-containing linear motifs, and represent the largest class of pTyr-recognition domains ^64^. SH2 domains have been studied biochemically and the peptide recognition specificity of different members of the SH2 family has been addressed by means of high-throughput approaches ^64–67^. These studies gave considerable insight in the understanding of SH2 recognition preference and in the SH2 mediated protein interaction network.

We have used MutateX to evaluate how post-translational modifications can modulate the interaction between SH2 domain and phosphorylated or non-phophorylated peptides, aiming at predicting the change of free energy of binding upon mutation of the SH2 peptide. This allowed us to understand which mutations influence the binding and thus correlate our results with the experimentally known binding specificity of phosphopeptides to SH2 domains.

We performed a MutateX saturation mutagenesis scan of different SH2 containing proteins complexed with phospho-tyrosine peptides∷CRK protooncogene, bound to a CRK-derived phosphopeptide (PDB ID 1JU5 ^68^), GRB2 SH2 domain bound to a phosphorylated peptide (PDB ID 1JYR ^69^) and SHP2 phosphatase complexed with a with RLNpYAQLWHR peptide (PDB ID 3TL0 ^70^. We selected these complexes because of their known heterogeneous binding specificities ^66,71^ and because 3D experimental structures were available.

We compared our results with the target consensus described in Tinti et al. ^66^, in which high-density peptide chip technology has been used to define the binding specificity of 70 different SH2 domains to pTyr peptides. Results from this article indicate that: i) CRK SH2 domain has a strong preference for Proline in position +3, ii) GRB2 SH2 domain has a strong preference for Asparagine in position +2 (from phospho-tyrosine) and iii) N-terminal SH2 domain of SHP2 has a preference for hydrophobic residues at positions –2, +1 and +3.

Following the MutateX run we analyzed the results by using the post-processing scripts from the MutateX suite of tools. At first, we used *ddg2distribution* to analyze the site averages of all the calculated ΔΔGs focusing our attention on the phosphopeptides. This tool was used to identify the residue positions that if mutated were impacting more on the free energy states of the complex. **(Fig. 6, center)**. Next, we used the *ddg2heatmap* tool to generate heatmaps of binding interface ΔΔGs, including all 20 natural occurring amino acids **(Fig. 6, left)**. This tool provided an immediate graphical representation of the residues, which were either more or less perturbing the interaction between the SH2 domains and the phosphopeptides, and gave more details on which substitutions are more likely to affect the interaction. We also used the *ddg2histo* tool to plot in graph bars the binding interface ΔΔGs for each residue in the phosphopeptides from positions −2 to +3 **(Fig. 6, right).**

**Figure 6.**
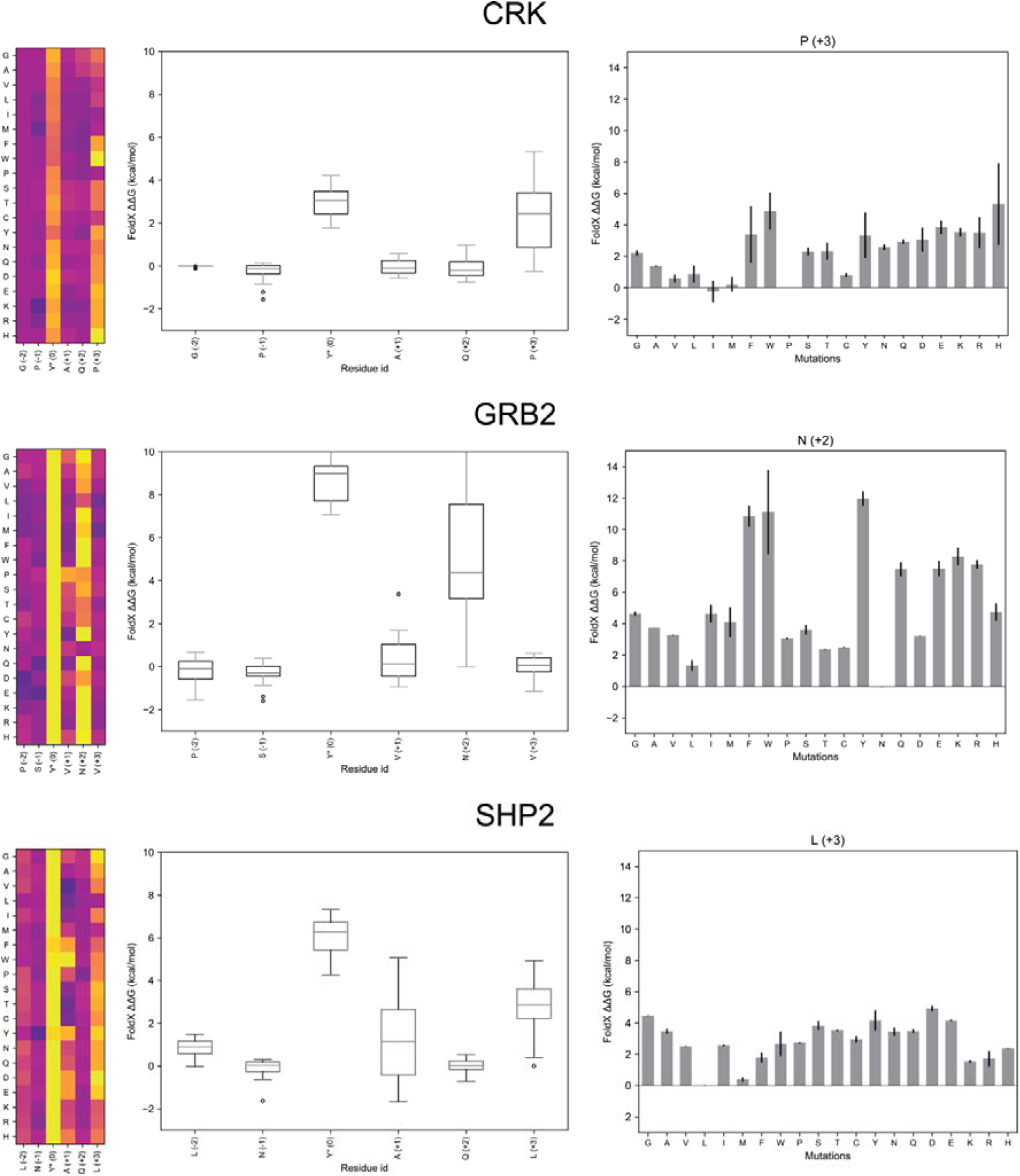
Changes in binding free energy in complexes of SH2-peptide upon phosphorylation. A) For each of the SH2 domain-complexes, we have plotted the calculated ΔΔG for the mutation of key residues of the phosphopeptide. These are shown as heatmaps, generated by ddg2heatmap, box plots, generated by ddg2distribution. The ΔΔG of a single key mutation sites are then shown more in details as histograms, generated using the ddg2histo tool.

As it is clearly shown in **Fig. 6**, in all the SH2/phosphopeptide complexes, mutations of the pTyr residues are associated with high ΔΔGs, indicating a strong impact on the binding affinity. The substitution tyrosine/pTyr shows elevated ΔΔGs, in agreement with experimental data according to which the tyrosine phosphorylation is prerequisite for the binding. Interestingly, the ΔΔGs were higher upon mutations in the residue positions that were shown to be mostly conserved in the different SH2 domain target consensus.

More in details the analysis showed that i) CRK SH2 domain displays a strong preference for Proline residue in position +3, while all the other mutations affect the interaction, except for branched-chain amino acids and Met, which present lower ΔΔGs; ii) GRB2 SH2 domain shows a strong preference in position +2 for Asn, with aromatic and charged amino acids being the most disrupting substitutions iii) SHP2 N terminal SH2 shows a moderate preference of hydrophobic residues in position −2 and +1 and a more pronounced preference in position +3 for Leu and Met.

Our predictions were consistent with experimental data which defined the SH2 domains consensus. Summarizing, this example clearly indicates that MutateX is a useful tool for the prediction of binding specificities and protein-peptide interaction analysis.

## Availability and Future Direction

MutateX is a collection of Python-based script and requires Python 2.7 and 3 or higher to be available on the system together with open source Python libraries, whose installation in straightforward on macOS and Linux distributions. MutateX comes with a standard installation script that takes care of installing all the necessary requirements as well. It is available at https://github.com/ELELAB/mutatex. The package also includes templates for input preparation, in-depth documentation and test cases. It should be noted however that in order to perform the mutagenesis calculation, the most recent version of the FoldX Suite needs to be available in the system. The FoldX Suite is free of charge for academic use and it is currently available at http://foldxsuite.crg.eu. While MutateX currently supports the FoldX Suite only, we plan on adding support for other software packages or web servers in the future, along with to provide customized support to other energy functions available in the literature and of potential interest for the user community.

The potential of MutateX is attested by recent publications where a preliminary suite of tools design by us was employed to perform saturation mutational scans for several systems of biological interest, such as enzymes of industrial and pharmacological interest ^72^, the Parkinson-related DJ1 protein ^73^, transcription factors as MZF1 ^74^, proteins involved in mismatch repair ^38,39^, and other disease-related targets ^38^. The aforementioned studies, along with the examples reported in this manuscript, provide insight into the mutational landscape of known cancer- or disease-related proteins, post-translational modifications and technologically important enzymes, allowing to predict the impact of these mutations in the context of important biological pathways or to design enzymes with different thermal stabilities.

MutateX provide a powerful complementary tool in experimental biochemistry, biotechnology and molecular/cellular biology, along with for the molecular understanding and classification of disease-causing mutations.

## Acknowledgments

The result of this research has been achieved using the DECI-PRACE15th HPC. The research has been also carried out thanks to the access to the Danish HPC Infrastructure Computerome. The research has been supported by LEO Foundation [LF17006], Carlsberg Distinguished Fellowship [CF18-0314] and Danmarks Grundforskningsfond [DNRF125]. Space limitation precludes the citations of many excellent works by colleagues in the fields discussed. We acknowledge these efforts here.

The authors are also extremely grateful to all the current and former Computational Biology Laboratory members at DCRC for extensive testing and usage of the software, which allowed us to optimize it. We also would like to thank Kresten Lindorff-Larsen, Amelie Stein and Rasmus Hartmann-Petersen for the interest and support that their groups have demonstrated for our project, and for having used the tool for their proteins of interest. We also would like to thank Kresten Lindorff-Larsen, Amelie Stein and Rasmus Hartmann-Petersen for their support and interest in MutateX. We are also grateful of the interest demonstrated by the FoldX developers in Luis Serrano’s group.

## Author contributions

Conceived and designed the project: MT, EP. Perform calculations and data visualization: TT (PTEN), MDM (PTEN), IP (p53), EM (SH2). Analyzed the data: TT (PTEN), MT (all), EP (all), IP (p53), EM (SH2). Contributed analysis tools, programming: TT, MT, TC, IP. Wrote the paper: MT and EP with inputs from all the coauthors.

